# Emergence of community behaviors in the gut microbiota upon drug treatment

**DOI:** 10.1101/2023.06.13.544832

**Authors:** Sarela Garcia-Santamarina, Michael Kuhn, Saravanan Devendran, Lisa Maier, Marja Driessen, André Mateus, Eleonora Mastrorilli, Ana Rita Brochado, Mikhail M. Savitski, Kiran R. Patil, Michael Zimmermann, Peer Bork, Athanasios Typas

**Affiliations:** European Molecular Biology Laboratory, Genome Biology, Heidelberg, Germany; European Molecular Biology Laboratory, Structural and Computational Biology, Heidelberg, Germany; Max-Delbrück-Center for Molecular Medicine, Berlin, Germany; Molecular Medicine Partnership Unit, Heidelberg, Germany; Department of Bioinformatics, Biocenter, University of Würzburg, Germany

**Keywords:** Microbiology, human-targeted drugs, growth inhibition, metabolomics, community resilience, drug biotransformation, drug bioaccumulation

## Abstract

Pharmaceuticals can directly inhibit the growth of gut bacteria, but the degree such interactions manifest in complex community settings is an open question. Here we compared the effects of 30 drugs on a 32-species synthetic community with their effects on each community member in isolation. While most individual drug–species interactions remained the same in the community context, communal behaviors emerged in 26% of all tested cases. Cross-protection, during which drug-sensitive species became protected in community, was 6-times more frequent than cross-sensitization, the converse phenomenon. Cross-protection decreased and cross-sensitization increased at higher drug concentrations, suggesting that the resilience of microbial communities can collapse when perturbations get stronger. By metabolically profiling drug-treated communities, we showed that both drug biotransformation and bioaccumulation contribute mechanistically to communal protection. As a proof-of-principle, we molecularly dissected a prominent case: species expressing specific nitroreductases degraded niclosamide, thereby protecting both themselves and sensitive community members.

## Introduction

Commonly prescribed therapeutics are associated with changes in the composition and function of the human gut microbiome ^1, 2^. Hundreds of drugs, including both antibiotics and those targeting human proteins, can directly inhibit the growth of commensal gut bacteria at physiologically-relevant concentrations ^3, 4^. Reciprocally, drug sequestration and/or biotransformation by gut bacteria can affect the bioavailability, efficacy, mode of action, and adverse effects of pharmaceuticals, thereby contributing to the interpersonal variability of drug responses ^5, 6^. Further molecular understanding of such drug–gut microbe interactions is crucial to design improved therapies with fewer side effects, including dysbiosis.

Several studies have used *in vitro*, *ex vivo*, and *in vivo* approaches to probe the impact of a limited set of drugs in diverse communities, showing both that drugs affect the community biomass and structure ^7–11^ and that communities affect drug activity, via mechanisms that include drug biotransformation ^6, 8, 12–14^ and bioaccumulation ^6^. Yet it remains unclear whether and to what extent such drug–microbial interactions in communities reflect the composite of effects observed in monocultures, and/or whether communal behaviors can mask or augment the single drug–microbe interactions. This understanding is crucial for our ability to predict responses of complex communities to drug treatment and to dissect drug-microbiota interactions based on simpler and controlled *in vitro* experimental setups.

Here, we assembled a synthetic community containing 32 representative species of the healthy human gut microbiota ^3^ and compared the effect of 30 diverse drugs on 21 species reproducibly detected in the community *versus* in isolation. We detected at least one species being protected or sensitized in the community setting for all drugs tested, and in total a quarter of all drug–microbe interactions (465/1823 cases) changing in the community setting. Cross-protection was the most frequent scenario, indicating that communities are more resilient to external insults than individual bacteria. However, such communal protective strategies decreased with increasing drug concentrations, while, cases of cross-sensitization increased. Thus, at higher drug concentrations, communities are disturbed the most – not only because more species may be targeted by the drug, but also because communities lose capacity for cross-protection and negative interactions (cross-sensitization) increase. Moreover, we demonstrated that both drug biotransformation and drug bioaccumulation contributed to many cross-protection instances, and mechanistically dissected a case of communal protection, identifying the species protecting the community and the enzymes degrading the drug. Using the knowledge about detoxifying species we could design synthetic communities that would enable growth of otherwise highly sensitive communities, opening the path for future use of such knowledge to optimize community composition to reduce adverse drug affects or increase drug efficacy. Overall, we provide insights into the degree of emerging behaviors upon treatment of microbial communities with drugs, identify some of their underlying mechanisms and map their dependence on drug concentration.

## Results

### Evaluating the impact of drugs on the composition of a complex synthetic community

We assembled a synthetic community of 32 species from 26 genera across 6 phyla (Suppl. Table 1), cultured at 37 °C under anaerobic conditions in Gifu Anaerobic Medium Broth, Modified (mGAM). Species were selected to be representative of the gut microbiome of healthy humans ^3^. From over 1,200 drugs previously tested against gut microbes in isolation ^3^, we selected 30 representative drugs that inhibited the growth of a variety of species and spanned many therapeutic areas (Figure S1A, Table S1). Out of the 30 tested drugs, 21 were human-targeted and 9 were anti-infective drugs (Figure S1A, Table S1). Drug concentrations were selected to always include 20 µM, as previously screened ^3^, and two additional concentrations, adjusted to the drug activity and colon concentrations (Figure S1A, Table S1). In almost all cases, concentrations were varied by factors of 4 (e.g. 5, 20, and 80 µM). Fifteen of the human-targeted drugs had at least one of the tested concentrations within a factor of three to the estimated colon concentrations (Figure S1A, Table S1). In contrast, antibiotics were tested in concentrations below those estimated to be present in the colon, since at such high concentrations many commensals were inhibited and the community barely grew *in vitro*.

To probe the effect of a drug on the community, we added the drug when combining the individual species that formed the community (Figure 1A). We passaged the community (1:50 dilution) once after 24 hours growth into fresh medium containing the respective drug, and grew the community for another 24 hours. At the end of the second passage, relative species abundance was determined using 16S amplicon sequencing. We found high correlations between the three biological replicates (median Spearman correlation between relative abundances r_s_ = 0.91 for controls and r_s_ = 0.89 for drug treatment, Figure S1B) and between the two technical replicates performed for each biological replicate (median r_s_ = 0.96 for controls and r_s_ = 0.97 for drug treatment, Figure S1B). To assess the impact of drug treatment on each species in the community, we normalized species abundances, using the community’s final optical density (OD) as a proxy for cell number (Figure 1B). To exclude that this step introduced any biases, we established that the main findings of this study also held when using only relative species abundances (Figure S1C, D). To determine the effect of drug treatment, we used the ratio of the species abundance between treatment conditions and untreated controls (Figure 1C). Eleven species were below the level of detection in untreated controls, and therefore we excluded them from further analysis. Interestingly, 7 of these low abundant species could be detected in at least one treatment condition (Table S1), indicating that drugs open niches for otherwise less fit community members.

**Figure 1:**
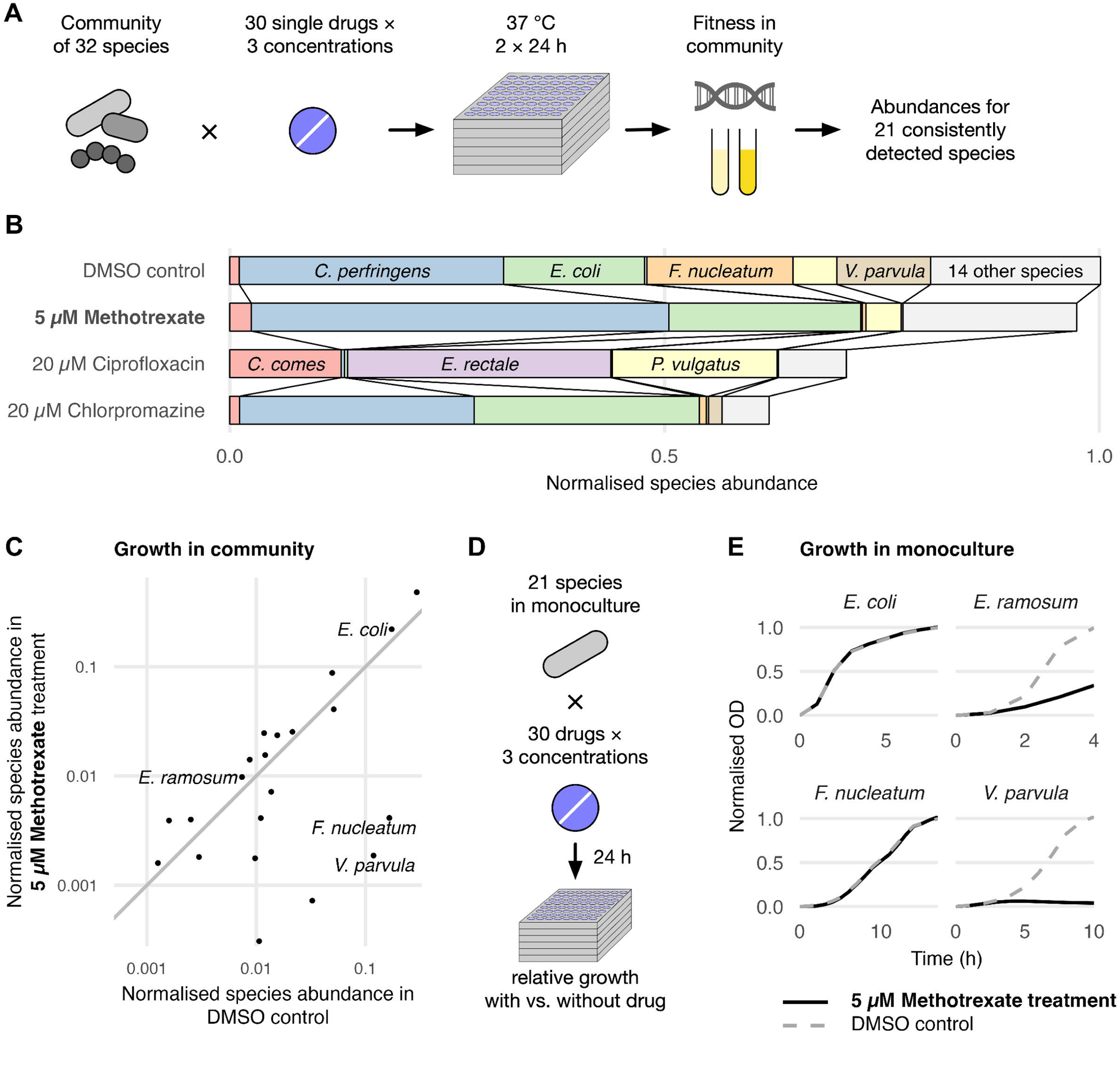
Measuring drug impact on gut bacteria in community and in isolation. (A) A community of 32 gut bacterial species was treated with 30 drugs in 3 concentrations. The community was immediately treated with drug upon assembly for 48 h, with an intermediate passage step (1:50 dilution) at 24 h. Community growth was measured by following OD, and relative species abundance by 16S rDNA amplicon sequencing at 48 h. Twenty one species were reproducibly detected in vehicle (DMSO control) and used thereafter. (B) Examples of species abundances in vehicle (DMSO control) and in three selected drugs and concentrations in the community. The abundance for each species of the community is normalized by the final OD of the community. (C) Example of the effect of 5 µM methotrexate in the community, comparing species normalized abundances between DMSO control and treated conditions. Species along the identity line were not influenced by the drug, while species below were inhibited in the presence of methotrexate. (D) The 21 species reproducibly detected in the community were also tested in the same panel of drugs as in A in isolation. Fitness was calculated by comparing the growth with *versus* without drug. (E) Examples of the effect of 5 µM methotrexate on selected species in monoculture compared to DMSO controls. While *E. coli* and *V. parvula* behaved the same in community (C) and isolation (E), *E. ramosum* and *F. nucleatum* were only inhibited in isolation (E) and in the community (C), respectively.

### Community behaviors emerge during drug treatment

All 21 species kept for further analysis were also treated individually with the 30 drugs (at 3 concentrations), and their growth was monitored (Figure 1D, E; Suppl. Table 1). We calculated the area under the growth curve (AUC) and normalized it for the vehicle (DMSO)-treated controls (Methods). For a total of 1823 drug–species combinations we compared the response to the drug in community and in monoculture; we excluded from this comparison 4 cases in which we did not obtain reproducible growth data in monocultures and the highest concentration for 3 drugs (chlorpromazine, ciprofloxacin, and doxycycline), in which the community did not grow at all. Three outcomes were identified (Figure 1C, E & 2A, using methotrexate as example): i) expected outcome-growth was similarly affected (*Veillonela parvula*) or unaffected (*Escherichia coli*) in both community and monoculture; ii) cross-sensitization (emergent communal behavior) - the species growth was not affected by the drug when alone, but its abundance was reduced in the community (*Fusobacterium nucleatum*); iii) cross-protection (emergent communal behavior) - the species was inhibited in monoculture but grew normally in community (*Erysipelatoclostridium ramosum*). To assess the degree of cross-protection and cross-sensitization per drug, we calculated the percentage of species that were either protected or sensitized in the community. For protection we divided by the total number of species that the drug inhibited in monoculture (as only those could be protected in community), whereas for sensitization we divided by the total number of species that grew normally in single-species experiments (Figure 2C, for the treatment amount closest to the gut concentration; Figure S2 for all treatment conditions). We observed at least one emergent behavior for all drugs probed, and in total 26% of drug-microbe interactions changed in the community setting. When taking into account only drug-sensitive species, protection in the community amounted to 47% of all cases, whereas community-specific sensitization was observed in 8% of all cases of resistant species. At the drug concentrations closest to the estimated human gut concentrations, these fractions were very similar with 49% and 9%, respectively. Overall, this suggests that numerous community-dependent protection and sensitization events can be expected upon drug treatment of human gut microbiotas, and those can vary between individuals, which harbor different community compositions.

**Figure 2:**
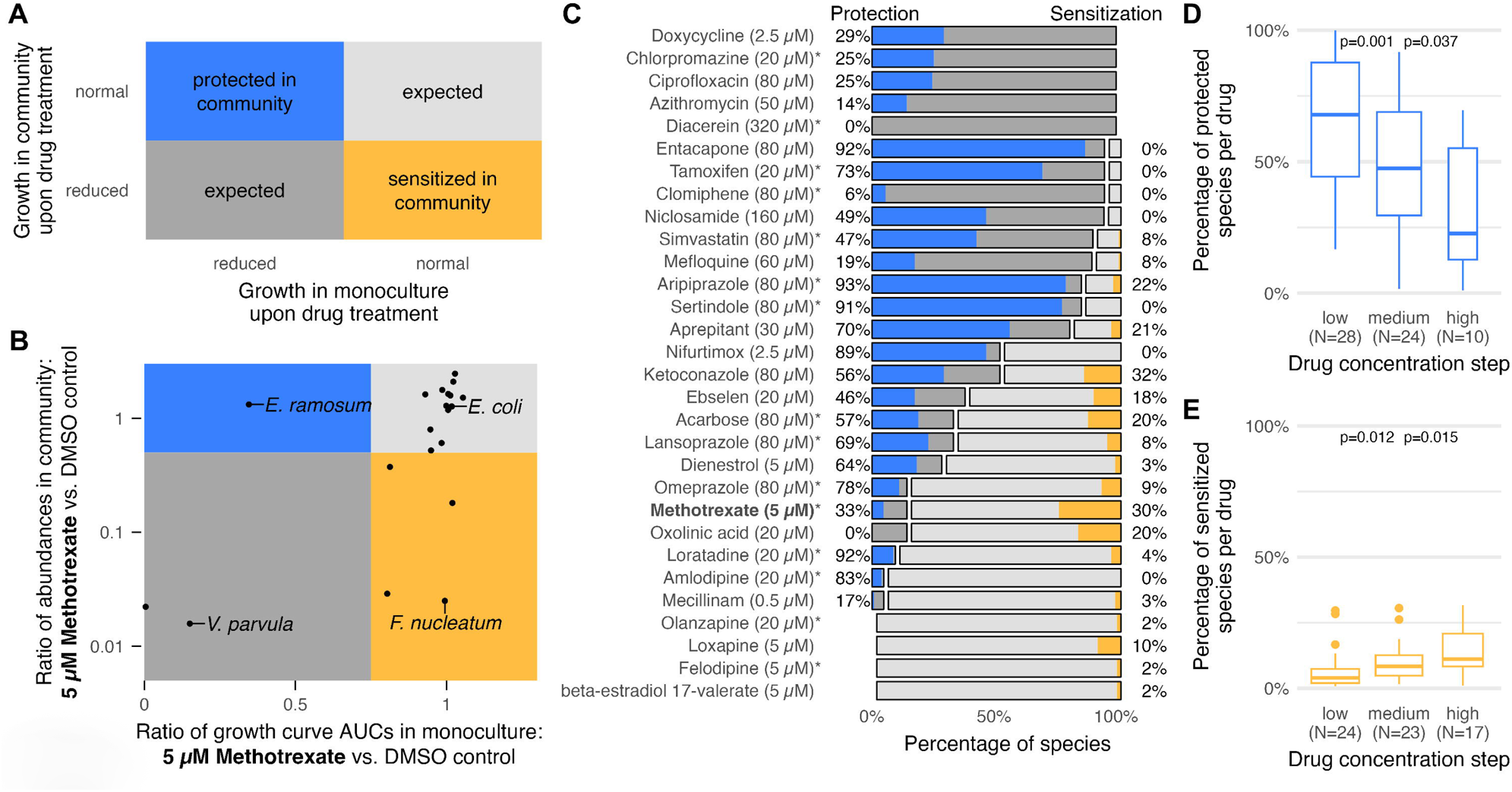
Emergent community behaviors are common upon drug treatment. (A) Comparing species growth in monoculture and in community in response to drug treatment. “Expected” refers to similar growth in both community and monoculture (light and dark grey areas), “protected in community” refers to species that are inhibited by the drug in monoculture, but remain relatively unperturbed in the community (blue area), and “sensitized in community” refers to species being unaffected in monoculture, but inhibited by the drug in the community (yellow area). (B) Examples of community emergent behaviors for 5 µM methotrexate treatment. *E. coli* growth was unperturbed both in monoculture and in the community (expected); *V. parvula* was inhibited in both cases (expected); *E. ramosum* was only inhibited in monoculture (protected in community); and *F. nucleatum* growth was only reduced in the community setting (sensitized in community). (C) Percentage of the 21 species across replicates that show expected or emergent behaviors (protection or sensitization in community) in all drugs tested at the concentration closest to the estimated colon concentration, or at 20 µM for those cases that the colon concentration could not be estimated (Figure S1). The asterisk denotes drug concentrations that are within a factor of three of the estimated intestinal concentration. (D-E) Higher drug concentrations bypass community resilience. A median of 68% sensitive species in 28 drugs were protected in the community at the lowest drug concentrations, but this significantly decreased to 47% (in 24 drugs) at the intermediate concentration, and to 23% (in 10 drugs) at the highest concentration (D). For drugs with resistant species (24 drugs) at the lowest concentrations sensitization occurred for a median of 4% of resistant species, and this increased to 8% (in 23 drugs) and 11% (in 17 drugs) at the intermediate and high concentrations, respectively (E). P-values were calculated using paired Wilcoxon signed-rank tests.

### High drug concentrations overwhelm community resilience

For each drug, we tested 3 concentrations. When starting from concentrations at which there was at least one sensitive species in monoculture (to be able to detect cross-protection), we could detect a significant drop in the percentage of protected species within the community across all drugs as the drug concentration increased (Figure 2D). *Vice versa*, the percentage of sensitized species significantly increased at higher drug concentrations (Figure 2E). Since the concentration steps were not always equally spaced, we also verified the concentration dependence in a separate model that uses the actual drug concentrations (Figure S3A-B). Overall, this means that the community stays relatively unaffected at low drug concentrations, since the perturbation is buffered and sensitive species are protected. In contrast, the impact on composition increases disproportionally at higher drug concentration, as not only the community fails to protect other members, but negative interactions emerge that sensitize otherwise resistant species.

### Bacterial drug biotransformation and bioaccumulation underpin emergent cross-protection

Many mechanisms could be driving the prevalent emergent behaviors we observed: interspecies interactions, new niches created by reduced growth of some species, altered stress responses in presence of others and/or modifications in the drug availability. Since previous work had showcased the extended ability of gut microbes to transform or intracellularly accumulate drugs ^6, 12, 13^, we decided to assess the degree to which emergent communal phenotypes, and especially cross-protection, could be explained by such phenomena. To this aim, we measured drug concentrations over time using liquid chromatography-coupled mass spectrometry (LC-MS) in the same synthetic community upon treatment with the same panel of drugs as above, typically at the concentration for which we observed the highest percentage of emergent behaviors (Figure S2). Samples were collected on the second day of treatment, at different timepoints after passage into fresh, drug-exposed medium (0, 1.25, 2.5, 5, 7.5 and 10 h after treatment). We collected two fractions per community: one containing the whole community (WC), i.e. both supernatant and bacteria, and one containing only supernatant (SN), to be able to distinguish between biotransformation and bioaccumulation ^6^. In parallel, we used only mGAM and the same time course to assess drug decay in the medium. For each time course, we normalized concentrations to the maximum value among the two first time-points of the time course and calculated AUCs. Overall biological replicates were consistent with mean standard deviation between replicates for all measurements being 8% for both community experiments and media control. From the time courses of each drug, we used AUCs to calculate the extent of biotransformation (media control minus WC), bioaccumulation (WC minus SN) and of both phenomena (media control minus SN) (Figure 3A and S4A). For example, the antibiotic ciprofloxacin was stable across all conditions; the proton-pump inhibitor lansoprazole decayed on its own, and this process was accelerated by the community; the antiparasitic niclosamide was rapidly metabolized by the community; and the antimalaria drug mefloquine was both bioaccumulated and biotransformed (Figure 3A).

**Figure 3:**
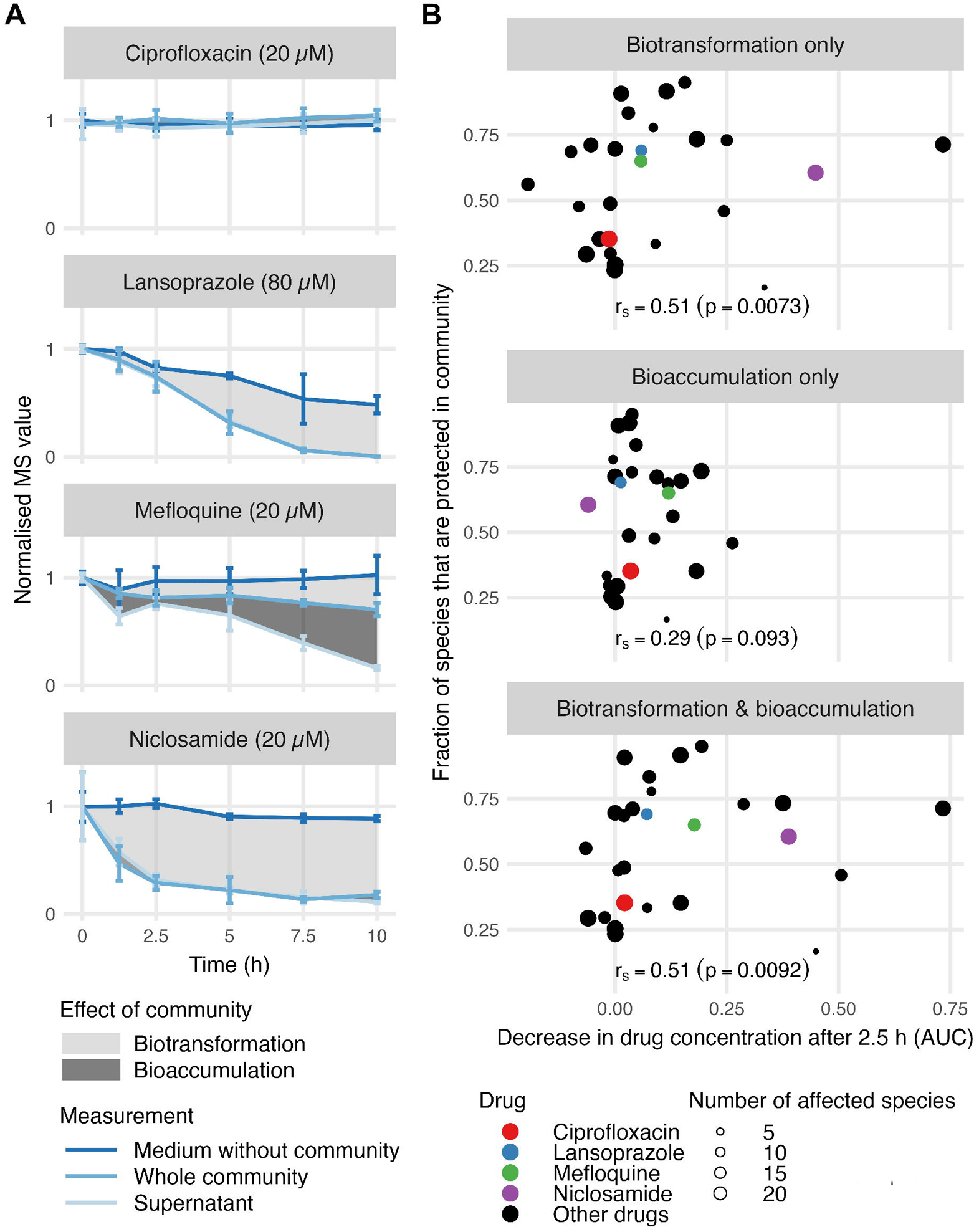
Bacterial drug biotransformation and bioaccumulation drive community cross-protection. (A) Representative examples of community effects on drug biotransformation and bioaccumulation during 10 h of time-course. Drug concentrations were measured by LC-MS in the 32-species synthetic community for 27 drugs (Figure S4A). For each time course, we normalized concentrations to the start of the time course (using time point with the highest raw measurement at 0 or 1.25 h) and calculated the mean drug concentration per time point. Bioaccumulation can be calculated by comparing the drug remaining in whole community and in supernatant (dark grey), and biotransformation by comparing the drug alone with the drug remaining whole community (light grey). Ciprofloxacin was stable in the community, lansoprazole was unstable in mGAM and decayed further in the community, mefloquine was biotransformed and bioaccumulated, and niclosamide was rapidly biotransformed by the community. (B) Fraction of protected strains correlated with degree of drug biotransformation and bioaccumulation by the community. P-values were calculated by permutation tests using 100,000 samples.

We found an overall positive correlation between the fraction of species that were protected by the community and the degree to which the drug was biotransformed by the community (Figure 3B), with the two earliest time points (1.25 and 2.5) showing significant correlations (Figure 3B and S4B). In contrast, bioaccumulation only mildly correlated with the fraction of both protected and sensitized species at early time points, with only the first timepoint showing a significant correlation with protection (Figure 3B and S4B-C). When combined with biotransformation, bioaccumulation often increased the overall correlation with the fraction of protected species in the community (Figure 3B and S4B). Overall, this means that the protective community effects can be at least partially explained by the drug being transformed (and to some degree accumulated) by one or more species in the community. Nonetheless, there are other mechanisms at play that remain to be elucidated in the future; for example, dienestrol was neither biotransformed nor bioaccumulated, yet 49% of susceptible species were protected in the community. In such cases, altered metabolism of drug-treated species may create new niches or neutralize the drug effect for the sensitive species. In contrast to communal protection, cross-sensitization did not correlate with biotransformation, and only partially correlated with bioaccumulation (Figure S4B-C). This could be due to the smaller number of cases identified. Having a better understanding of what drives cross-protection, we decided to study some of the underlying mechanisms in further detail.

### Different species protect against different nitroaromatic drugs in communities

All three nitroaromatic drugs used in the screen, the antiparasitic drugs niclosamide and nifurtimox, and a drug to treat Parkinson’s disease, entacapone, were rapidly biotransformed by the community (Figure S4A). At the highest tested concentration, these drugs inhibited on average 92% of the tested species in monoculture (Table S1). In line with the observed biotransformation, we found 47% of these sensitive species to be protected by the community (Figure S2). Since both activation and detoxification of nitroaromatic compounds rely on reduction of the nitro group ^15, 16^, we wondered whether it is the same set of microbes with potent nitroreductases that efficiently transformed the drugs to an inactive form.

To explore this further we selected seven species from the community, which covered a wide range of sensitivities to niclosamide in monoculture (Figure 4A). Using LC-MS, we checked for the ability of these seven species to reduce niclosamide to its amine form. Three out of the four (partially) resistant species in monoculture (*Roseburia intestinalis*, *Coprococcus comes* and *Fusobacterium nucleatum*, Figure 4A) stoichiometrically and rapidly reduced niclosamide to aminoniclosamide (Figure 4B-C and S5A), which was not toxic to any of the gut bacterial isolates tested (Figure S5B). Indeed, this biotransformation of the drug to the inactive amino form resulted in protection of niclosamide-sensitive species from niclosamide toxicity, as we showed by growing sensitive species in spent media of bacteria with biotransformation capacity (Figure 4B). The degree of protection of sensitive species was directly related to the ability of the protecting species to degrade the drug (Figure 4C).

**Figure 4:**
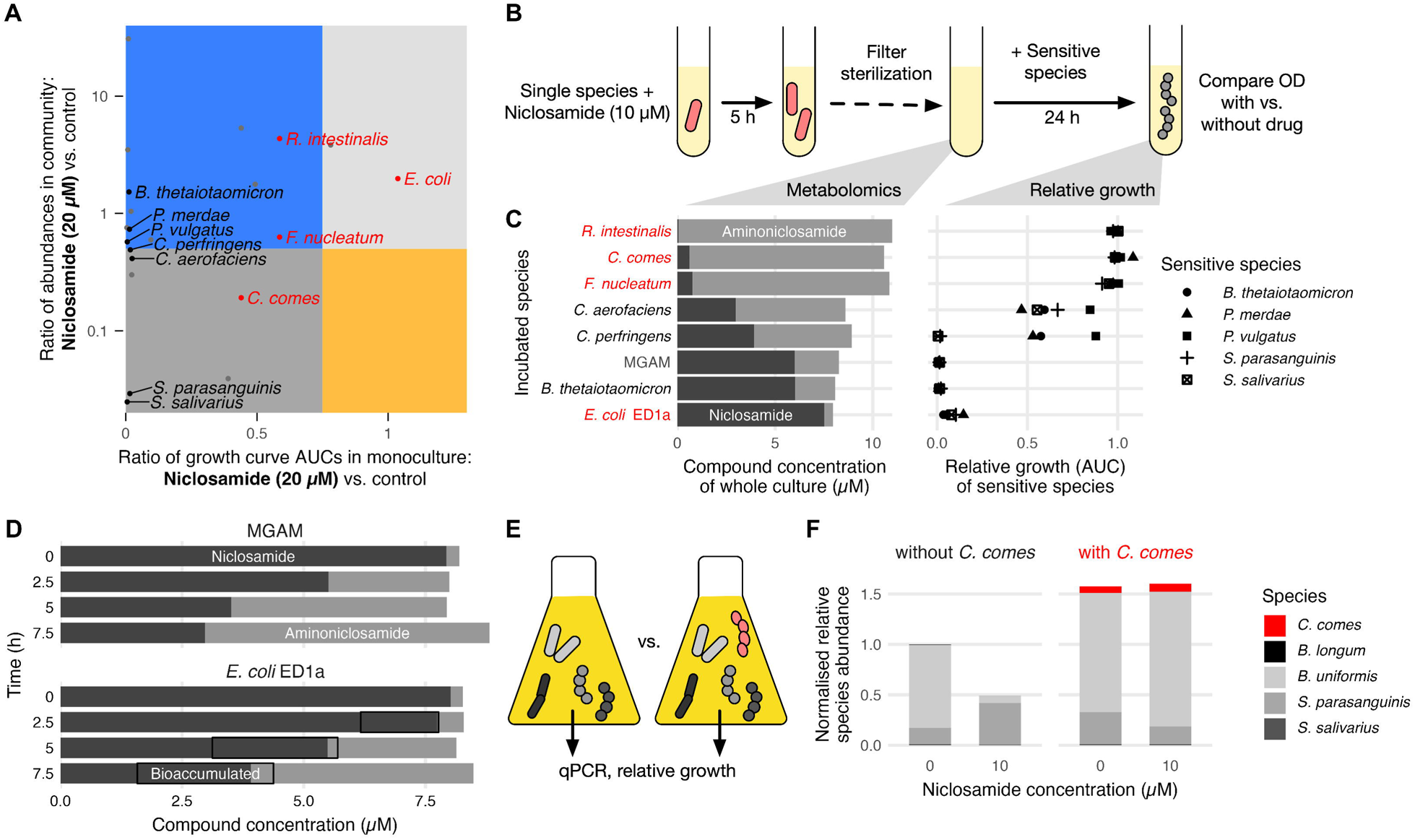
Specific species protect the community from niclosamide. (A) Community-emergent phenotypes in 20 µM niclosamide. Representation of species that grow in community as expected from monoculture (grey areas), that are cross-protected (blue area) or cross-sensitized (yellow area) in the community. Species used in further tests are highlighted: partially/full resistant species to niclosamide in red and sensitive species in black. (B) Experimental strategy used to disentangle contributions of individual species to community protection from niclosamide. Species were incubated with niclosamide for 5h, after which cultures were analyzed by targeted metabolomics to quantify niclosamide and aminoniclosamide. In separate experiments, the same species were treated with vehicle (DMSO control) or 10 µM niclosamide for 5 h, after which cultures were filter-sterilized. Spent media from these cultures was mixed 1:1 with fresh mGAM, and used to grow other niclosamide-sensitive species. Growth of these other species in spent media from drug treated cultures was normalized to their growth in spent media from DMSO cultures (controls). (C) Left panel, following the scheme in (B), quantification of niclosamide and aminoniclosamide in culture media of the indicated species (text color as in A) after 5h incubation with niclosamide. Right panel, relative growth of the indicated species (sensitive to niclosamide – see A) in spent media from the niclosamide treated cultures. (D) Quantification of niclosamide and aminoniclosamide concentrations in medium control (upper panel) and in cultures of *E. coli* ED1a cells after treatment with 10 µM niclosamide for the indicated times (lower panel). Boxes denote the amount of bioaccumulated drug in *E. coli* ED1a. To determine them, we subtracted the measured compound concentrations in supernatants from those measured in whole cell cultures. Niclosamide is more stable (i.e. less reduced to aminoniclosamide) in *E. coli* ED1a cell cultures (lower panel) than in the medium control (upper panel), because it is bioaccumulated in ED1a cells. (E-F) Addition of a niclosamide-detoxifying species can rescue a synthetic community from niclosamide treatment. *C. comes* restores growth of a niclosamide treated community, yielding species abundances similar to that of the untreated control. In addition to drug protection, *C. comes* promotes the overall community growth via unknown mechanisms.

In contrast to the other three (partially) resistant species, we found that the most resistant species, *E. coli* ED1a, could not degrade niclosamide, and the drug was even more stable in the culture than in the medium control (Figure 4C). We hypothesized that *E. coli* ED1A bioaccumulates niclosamide, preventing the non-enzymatic reduction of the drug ^17^ in the medium. Indeed, by using drug-treated cultures and spent media, we showed that a significant amount of niclosamide bioaccumulated in *E. coli* ED1a (∼ 2 µM in 5 h, Fig. 4D) without affecting the fitness of the organism. However, this amount of bioaccumulation in *E. coli* ED1a allowed only limited protection of other sensitive species in the spent media (Figure 4C).

In stark contrast to its inability to reduce niclosamide, *E. coli* ED1a had high protective capacity for nifurtimox, completely transforming 20 µM nifurtimox in 5 h, and fully protecting nifurtimox-sensitive species (Figure S5C-D). Another fully resistant species to nifurtimox, *Streptococcus parasanguinis* (Figure S5C), only partially metabolized the drug after 7.5 h incubation and hence offer limited protection to sensitive species (Figure S5D). Overall, this highlights the diversity of drug-microbe interaction mechanisms, even with drugs with the same functional group that is subject to the same type of bacterial transformation. It also highlights that the degree of resistance is not predictive of the ability to protect other species in the community, as resistance mechanisms differ.

We further wondered whether we could use this knowledge of selective protection to engineer communities that are more robust to drug treatment. To do this, we grew a small community of gut bacteria sensitive to niclosamide, composed of *Bifidobacterium longum*, *Bacteroides uniformis*, *S. parasanguinis*, *S. salivarius* (Minimal Inhibitory Concentrations ≤ 1.25 µM), to which we added or not the detoxifier *C. comes* (Figure 4E)*. C. comes* restored the growth of the community, even after 10 µM niclosamide treatment, yielding species relative abundances similar to those of the untreated control (Figure 4F). *C. comes* provided communal resistance to niclosamide, despite only growing modestly in the community (Figure 4F), highlighting that protection can be also offered from members that are low abundant in the community.

### Nitroreductase expression and specificity determine ability to reduce nitroaromatics

To pinpoint the mechanism by which niclosamide cross-protection occurred, we looked for the nitroreductases encoded in the genomes of the protecting and sensitive species. Oxygen insensitive or type I pyridine nucleotide-flavin mononucleotide [NAD(P)H/FMN]-dependent nitroreductases reduce nitroaromatics through stepwise additions of two-electrons to nitroso-, hydroxylamino- or amino-aromatics ^18^. FMN-dependent nitroreductases constitute a large and diverse family of proteins, mainly present in bacteria, which have been recently reclassified into 14 subgroups according to their sequence similarities ^19^. We found that all species in our community encode at least two of these enzymes belonging to different subgroups (Table S2). To further understand the basis of niclosamide reduction, we focused on two species: a) *R. intestinalis*, moderately resistant to niclosamide and a good protector, encoding only two putative FMN-dependent nitroreductases (other resistant species encoded more nitroreductases), and b) *P. vulgatus*, sensitive to niclosamide (MIC = 0.625 µM), but intriguingly encoding seven putative nitroreductases (Table S2). To test the ability of each of these nitroreductases to degrade niclosamide, we cloned and overexpressed them in the sensitive *P. vulgatus*. Overexpressing any of the two *R. intestinalis* nitroreductases conferred 8-fold or more increase in MIC to niclosamide for *P. vulgatus* (Figure 5A & S5E), suggesting that these enzymes are responsible for the resistance of *R. intestinalis* in niclosamide. Indeed, nitroreductase C7GA87 was highly expressed in *R. intestinalis* (Figure 5B). Interestingly, overexpression of two out of the seven *P. vulgatus* nitroreductases conferred also significant resistance to niclosamide (Figure 5A & S5E). We reasoned that these proteins should be silent or lowly expressed from endogenous locus, and hence *P. vulgatus* is sensitive to niclosamide. Indeed, both proteins had low abundance, which was not further induced by niclosamide in monoculture (Figure 5B). Overexpression of nitroreductase Pv2039 led to > 16-fold increase in protein levels (Table S2) and 32-fold higher niclosamide resistance (Figure 5A-B). As expected, only *P. vulgatus* overexpressing the *R. intestinalis* nitroreductases or the endogenous Pv2039 allowed other niclosamide-sensitive strains to grow (Figure 5C-D).

**Figure 5:**
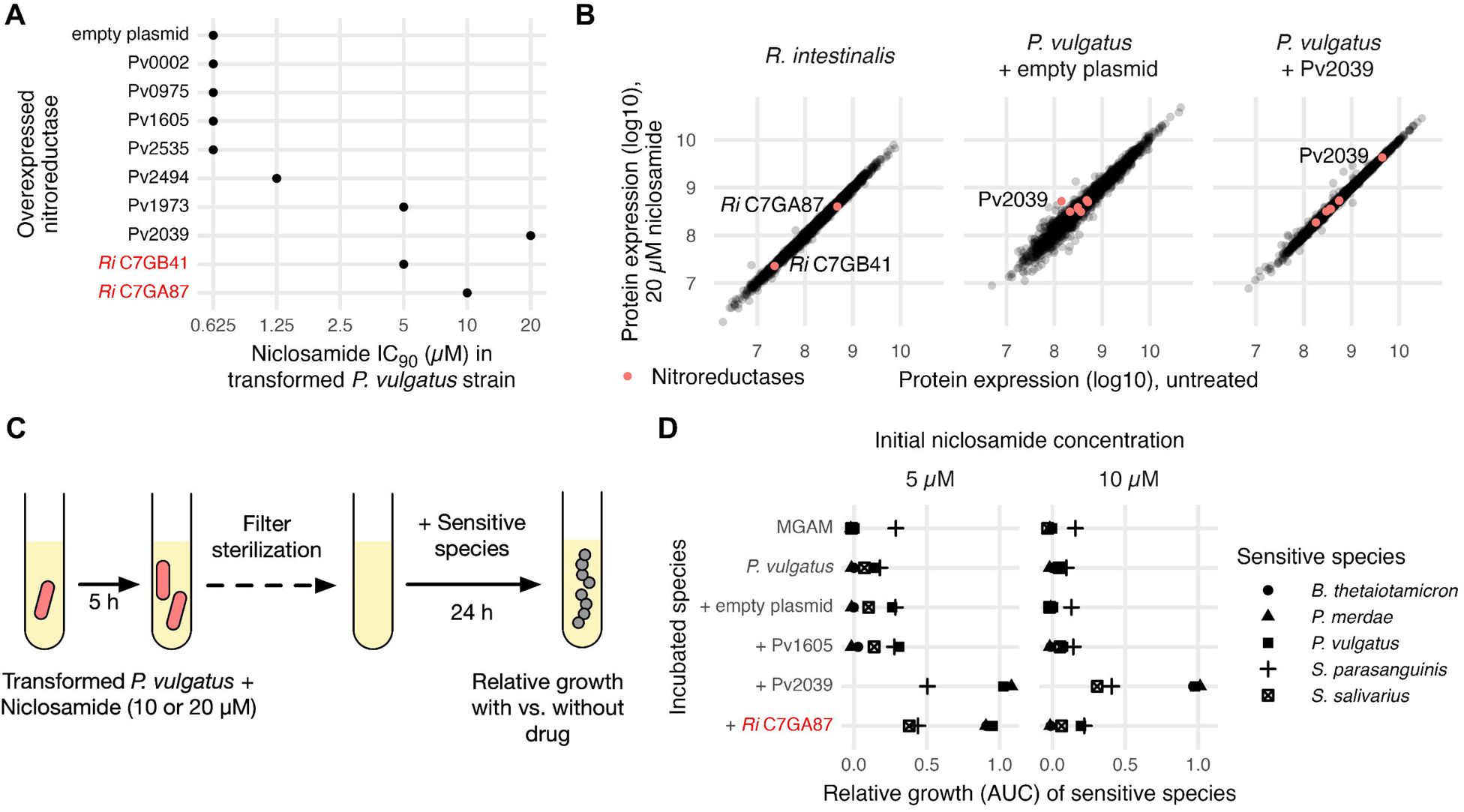
Specific nitroreductases protect the community from niclosamide. (A) Effect of heterologous nitroreductase overexpression on *P. vulgatus* IC_90_ (µM) to niclosamide. *P. vulgatus* strains transformed with empty plasmid, or plasmids overexpressing different *P. vulgatus* or *R. intestinalis* nitroreductases, as indicated, were treated with niclosamide, and growth was monitored by measuring the OD_578_ during 24 h. IC_90_ represents the concentration at which 90% of the growth is inhibited (see Figure S5E for IC curves). (B) Comparison of proteome expression between niclosamide (20 µM for 30 min; y-axis) and vehicle (DMSO control, x-axis) treated cultures of *R. intestinalis*, *P. vulgatus* transformed with an empty plasmid, and *P. vulgatus* over-expressing the nitroreductase Pvu2039. The two putative *R. intestinalis* nitroreductases and the seven putative *P. vulgatus* nitroreductases are highlighted in red. (C) *P. vulgatus* expressing different nitroreductases has different abilities to protect niclosamide sensitive species. Similar to experiment in Figure 4B-C, *P. vulgatus* strains carrying different nitroreductases were treated with DMSO control, 10 µM, or 20 µM niclosamide for 5 h, after which cultures were filter-sterilized. Spent media from these cultures was mixed 1:1 with fresh mGAM, and used to grow niclosamide sensitive species. (D) Following the scheme in (C), growth of the indicated strains in spent media from niclosamide treated cultures. Growth of the sensitive strains in spent media of treated cultures was normalized with their growth in spent media from untreated cultures (controls).

Overall, these results indicate that nitroreductases are specific to the substrate. When we overexpressed the same nitroreductases as the ones above in *P. vulgatus*, resistance to nifurtimox did not change (Figure S5E), which was consistent with the sensitivity of both *P. vulgatus* and *R. intestinalis* to nifurtimox (Figure S5C). Overall, our data suggest that that many species may have the capacity to biotransform drugs, but the respective selective enzymes may not be expressed and/or induced upon drug treatment.

## Discussion

In recent years, advances in gut microbiota culture techniques have made it possible to systematically determine the direct interactions between hundreds of commonly used drugs and specific members of the gut microbiota ^3, 7, 13^. However, little is known about whether these direct drug-bacterial species interactions are relevant when the same strain/species is part of a bacterial community. Here we show that 74% of a total of 1823 directly determined drug–species interactions remained the same in the context of a community. Nevertheless, communal behaviors were substantial, and were present in every drug we tested. Protection of sensitive species was the most common outcome, and even more prevalent at low drug concentrations. At higher concentrations, closer to that found in the colon, community protection decreased, with more species behaving the same as when growing alone. Hence, despite the occurrence of communal behaviors, single species–drug interactions are relatively good predictors of what will happen to a species when it is part of a community. This is consistent with decades of clinical work where antibiotic sensitivity of enteric pathogens is tested in isolation and not in community settings.

In other contexts, it has been shown that positive interactions between community members increase with the degree of species dissimilarity ^20^. We assembled a diverse community of 32 bacterial species belonging to 26 genera, representative of the human intestinal microbiota from healthy individuals ^3, 21^. It would be interesting to establish whether communities with less phylogenetic diversity, typically found in dysbiosis, maintain the same degree of resilience and cross-protection to drugs as that observed here.

Based on targeted metabolomics data, we established that bacterial drug biotransformation partially explains community protection phenotypes. This implies that in most cases drug biotransformation yields harmless or less toxic products. In our *in vitro* setting, drug bioaccumulation only mildly correlated with protective phenotypes. This could be because we added the drug during passaging, meaning that at the beginning there is a relatively low bacterial biomass that could accumulate the drug. Bioaccumulation has only been recently reported for drugs and gut microbes ^6^. In addition to the previously reported drugs, we found here that intestinal bacteria bioaccumulate four more drugs: ebselen, mefloquine, simvastatin and tamoxifen. Ebselen may accumulate due to its ability to bind to thiol groups^22^. In the case of tamoxifen or mefloquine, bioaccumulation could be a result of their ability to interact with bacterial membranes, as previously shown for *Bacillus stearothermophilus* and *E. coli*, respectively ^23, 24^. The case of simvastatin is interesting, since previous studies suggest a link between microbiome composition and the efficacy of the drug in lowering LDL cholesterol in patients ^25–29^. Statins are one of the most prescribed drug groups in western countries ^30^, and hence commonly found into wastewater ^31^. Due to their high water-octanol partition coefficient, statins tend to bioaccumulate in aquatic animals, causing a serious environmental problem ^32, 33^. It would interesting investigate in the future if bioaccumulation of statins in bacteria has implications for drug efficacy in patients, and/or may also affect the accumulation of the drug in environmental reservoirs.

Our data underline the broad ability of the microbiota to transform xenobiotics. Working with a bottom-up assembled synthetic community allowed us to gain insights into the biotransformation mechanisms that lead to communal protection. We focused on nitroaromatic compounds, and observed that the nitroreductases that reduce them are rather specific. For example, both nitroreductases of the niclosamide-resistant *R. intestinalis* efficiently reduced niclosamide but not nifurtimox. Importantly, *P. vulgatus* encoded nitroreductases that could render it resistant to niclosamide, but did not express them in relevant amounts even in the presence of the drug. This denotes that gut bacteria likely have an even larger potential to transform drugs than previously appreciated, which would render them able to evolve resistance if exposed to the drug for longer periods.

Albeit less frequent at low drug concentrations, cross-sensitization became more dominant as emergent behavior at higher drug concentrations. Cross-sensitization may arise from altered biotransformation capacities of the strain accumulating the drug ^6^ or induction of toxic stress responses by strains targeted by the drug. Although the underlying mechanisms behind such cases remain to be explored, cross-sensitization further disturbs community composition. Together with loss of cross-protection at high drug concentrations, they presumably lead to an accumulative strong impact in community stability.

In summary, we established that most drug–gut bacteria interactions remain the same in communities, but cross-protection and/or cross-sensitization of specific community members happens for almost every drug. The knowledge of such interactions that change in the community setting could help to build more accurate predictive models of community responses to drug in the future. We further showed that communities have higher resilience than individual species to certain drugs, but up to a certain concentration limit, after which communal protection drops and cross-sensitization and individualized behaviors prevail. This resilience is partially explained by bacterial drug biotransformation activities, which is facilitated by the expanded functional diversity of the community. However, this is not the only mechanism by which communities become resilient. Understanding the remaining underlying mechanisms can facilitate the targeted design of designed communities that are resilient to specific drugs, as we show here for niclosamide. Our work also contributes to expanding the notion that drug bioaccumulation is widespread among gut bacteria ^6^. Understanding the drivers of bioaccumulation, as well as its potential to affect drug mode of action, merit both deeper exploration. Overall, we have used a bottom-up approach to assess the degree of communal behaviors that emerge in response to drugs, and to map some of the underlying mechanisms that govern interactions between drugs and gut bacteria.

## Acknowledgments

We thank Justin Sonnenburg and Carlos GP Voogdt for plasmids; Jacob Bobonis for providing the *E. coli* conjugation donor; Ece Kartal for help with primers design for qPCR; the EMBL Gene Core Facility for services and experimental assistance; Thomas Sebastian B Schmidt and Georg Zeller for analysis and feedback of pilot experiments; and Pascale Cossart and Carlos GP Voogdt for feedback on the manuscript. We acknowledge EMBL, ERC grant uCARE (ID 819454), and the H-2020 Twinning project: SymbNET Genomics & Metabolomics in a Host-Microbe Symbiosis Network (Grant: 952537) for funding. S.G.-S. and L.M. were supported by the EMBL Interdisciplinary Postdoc programme under the Marie Sklodowska Curie Actions COFUND (grant numbers 664726 and 291772, respectively).

## Author contributions

This study was conceived by K.R.P., P.B., and A.T; designed by S.G.-S., M.K., L.M., M.Z., and A.T. M.K. pre-processed, curated, analyzed and/or contributed to the analysis of all experimental data. S.G.-S., L.M., and M.D. performed community experiments; S.G.-S. and A.R.B. performed high-throughput DNA extraction; S.G.-S., S.D., and E.M. performed metabolomics experiments and analyzed metabolomics data; S.G.-S. and A.M. performed proteomics experiments and analyzed proteomics data; S.G.-S. performed niclosamide and nifurtimox experiments; M.S., M.Z. and A.T. supervised experimental work. Data interpretation was performed by S.G.-S, M.K., K.R.P., M.Z., P.B., and A.T. S.G.-S., M.K., and A.T. wrote the manuscript with feedback from all authors. M.K. and S.G.-S. designed the figures with input from K.R.P., M.Z., P.B., and A.T.

## Declaration of interests

The authors declare no competing interests.

## Methods

### Bacterial species and growth conditions

Species used in this study were purchased from DSMZ, BEI Resources, ATCC and Dupont Health & Nutrition, or were gifts from the Denamur Laboratory (INSERM) and processed as previously described ^3^. All species (monoculture or community) were grown in mGAM (HyServe GmbH & Co.KG, Germany) except the monocultures of *Veillonella parvula* and of *Bilophila wadsworthia*, which were grown in Todd-Hewitt Broth supplemented with 0.6% sodium lactate and mGAM supplemented with 60 mM sodium formate and 10 mM taurine, respectively. Media was pre-reduced to a minimum of 24 h under anoxic conditions (2% H_2_, 12% CO_2_, 86% N_2_) in an anaerobic chamber (Coy Laboratory Products Inc.). Unless otherwise specified, experiments were performed at 37°C and under anaerobiosis, species were inoculated from frozen stocks into liquid culturing media, and passaged twice overnight to ensure robust growth, and drug stocks in DMSO were pre-reduced overnight in the anaerobic chamber inside an ice-box at 4 °C.

A representative core of species in the human gut microbiome were selected as previously described ^3, 21^. From this core, 32 diverse species, differing more than 3% on their 16S rDNA sequences at the V4 region, were selected for the screening.

### Chemical screening of a bottom-up assembled bacterial community of 32 species

#### Screening plates preparation

Drugs were dissolved with the appropriate solvent (Table S2) at a concentration 100-fold higher than the screening concentration, distributed in 96-well V-bottom plates (Greiner, 651261), each well containing 11 µl of dissolved compound or vehicle, and stored at -30°C for up to 1 month. One day before the experiment, drug plates were thawed, and each of the 11 µl of drug or vehicle per well were added to 539 µl/well of mGAM in 96-deep well plates (Costar 3959) using the Biomek FXP (Beckman Coulter) liquid handling system. These plates were pre-reduced in the anaerobic chamber overnight (“Community Plates Day1”). Inoculation of community passage 1. For community assembly, the optical density (OD) was individually measured at 578 nm for the 32 species. These were added together into 200 ml of mGAM with the volume required to reach a total of 2x the desired initial OD of 0.01: therefore, each species was added at an OD of 0.0006. 550 µl of the assembled community was added into each well of the “Community Plates Day 1” with an epMotion 96 (Bio-Rad) semi-automated electronic 96 channel pipette. Final drug concentrations are described in Table S1 and each well contained 1% DMSO. After inoculation, 100 µl/well were transferred from the “Community Plates Day 1” to U-bottom shallow 96-well plates (“Community Growth Plates Day 1”) (Fisher Scientific 168136), sealed with breathable membranes (Breathe-Easy, Sigma), and incubated at 37°C. These plates were used for monitoring community growth after drug treatment, for which OD_578_ was measured with a microplate spectrophotometer (EON, Biotek) every hour during 24 h, with shaking only few seconds before OD measurement. The “Community Plates Day 1” were also sealed with a breathable membrane (AeraSeal, Sigma) and incubated during 24 h. The untreated community grew to an average OD_578_ of 1 (7 generations). Inoculation of community passage 2. After the initial 24 h incubation, the drug-treated community was passaged with a 1:50 dilution (OD_578_ of 0.02 for the untreated community) into a new drug-containing deep-well plate for an additional 24 h incubation as follows: during the initial 24 h incubation period, new drug plates were thawed, and the 11 µl of drug or vehicle was added to 1067 µl/well of mGAM in 96-deep well plates (“Community Plates Day 2”). These were pre-reduced in the anaerobic chamber overnight. Wells in the “Community Plates Day 1” were mixed with the 96 channel epMotion pipette, and from here 22 µl were transferred to the “Community Plates Day 2”. Of these, 100 µl were transferred to 96-well U-bottom plates (“Community Growth Plates Day 2”), for growth curve acquisition, as described above. The “Community Plates Day 2” were incubated for 24 h anaerobically and at 37°C. The untreated community grew to an average OD_578_ of 1 (6 generations). After 24 h of incubation, cell pellets were collected by centrifugation. DNA extraction and 16S rDNA sequencing were performed as described below. These experiments were performed in 3 biological replicates with 2 technical replicates each.

#### Genomic DNA isolation

Genomic DNA isolation was done as previously described ^4^. Briefly, DNA was extracted in a 96-well plate format using a Biomek FXP (Beckman Coulter) liquid handling system or in single tubes depending on the number of samples. Cells were first washed with PBS and resuspended in 281 µl of cell suspension solution (MP GNOME DNA kit). Cell suspension was treated with lysozyme (25 µl; 400.000 U/ml) and incubated for 1 h at 37 °C. Cell suspensions were then further lysed by three freeze/thaw cycles using liquid nitrogen, before the addition of 15.2 µl of cell lysis solution (MP GNOME DNA kit) and 20 µl of RNA mix (MP GNOME DNA kit). A last step of lysis was performed using glass beads (Glasperlen, Edmund Bühler) by bead beating twice for 5 min at 30 Hz in a Tissue Lyzer II (QIAGEN). Lysates were then incubated for 30 min at 37 °C with shaking. 12.8 µl of protease mix (MP GNOME DNA kit) was subsequently added and the lysates were incubated for 2 h at 55 °C. After a 5-min centrifugation step at 3.200 x g, 200 µl of supernatants were collected and mixed with 100 µl of TENP buffer ^34^ (buffer: 50 mM Tris-HCl, pH=8, 20 mM EDTA, 100 mM NaCl, 1% w/vol polyvinlylpolypyrrolidone), and with 75 µl salt out solution (MP GNOME DNA kit). These were incubated for 10 min with at 4 °C. After a 10 min centrifugation at 3200 x g, 200 µl of supernatant were transferred to a clean plate. 500 µl of ice-cold ethanol and 70 µl of 3M NaOAc pH 5.2 were added. The solution was kept at -30 °C overnight. The next day, the plates were centrifuged at 4 °C, 3.200 x g for 45 min. The supernatant was carefully removed and the pellets were washed with 400 µl of ice-cold 70% ethanol. After a 20 min centrifugation at 3200 x g at 4 °C, all the supernatant was removed and plates were dried in a chemical hood for 30 min. DNA was resuspended in 70 µl water overnight at 4 °C.

#### 16S rDNA sequencing

The 16S libraries were then prepared for sequencing using a two-step PCR method according to ^35^, using the Phire Hot Start II DNA polymerase (Thermo Scientific). Briefly, the V4 region was amplified by a first PCR with the 515F/806R primers (Table S2). The resulting amplicons were subsequently amplified again using barcoded primers that contain Illumina adaptors. These libraries were sequenced in the EMBL GeneCore sequencing facility on an Illumina MiSeq (250 base pairs, paired-end).

#### Processing of 16S data and quantification of species abundances

To estimate the species abundance, a database of 16S rRNA regions was constructed by manually querying the SILVA rRNA database ^36^ and extracting the representative sequence from each of our 32 species. Amplicon sequencing reads were then mapped against this database using MAPseq v1.2. ^37^. Paired reads were mapped independently and assignments were only considered upon agreement. Abundance estimates were then produced by counting the number of reads mapping to each genome included in the study. Eleven species whose median read count in controls was below 10 were designated as rare species and excluded from the subsequent analysis, as for these species abundance ratios would be unreliable. Relative species abundances were calculated by dividing the number of reads mapping to each species by the total number of reads for a sample. To estimate absolute species abundances, we multiplied the relative abundances by the final normalised OD (which was set to 1 for control wells). To ensure that relying on OD as a proxy for cell counts does not lead to large distortions, we also calculated standardized relative abundances by dividing the relative abundances within each condition by the 75^th^ percentile of the relative abundances. We chose the 75^th^ percentile instead of the median, as this is more robust to the case where the growth of many species is inhibited.

#### Quantification of treatment effects

Control abundances were separately determined for each biological and technical replicate by calculating the mean abundance across six control wells. For each species and treatment condition, we calculated the ratio of treatment and control abundances to estimate the effect of drug treatment on each species independently of its absolute abundance. Species whose abundance was decreased to 50% or less upon drug treatment were designated as being reduced in the community.

#### Community metabolomics

LC-MS analyses were employed to assess the presence/absence of a drug after incubation with the community of 30 species (compared to the initial community, Ruminococcus bromii, and Ruminococcus torques, which did not grow reliably in the community, were not added as cells did not grow from stocks). Drug plates, community assembly, passages and incubations were performed as described above, with the only exception that the community inoculation in the 2^nd^ passage was performed in a final volume of 1.5 ml per well. Final drug concentrations are listed in Table S1. Immediately after inoculating the community in the 2^nd^ passage: i) 100 µl were transferred to 96-well U-bottom plates for growth curve acquisition, as described above, ii) 100 µl were transferred to 96-well shallow U-bottom plates and immediately stored frozen at -80 °C (0 h time point), and iii) 150 µl were transferred to 5 different 96-well shallow U-bottom plates, which were incubated anaerobically at 37 °C for 1.15 h, 2.5 h, 5 h, 7.5 h, and 10 h, respectively. At each time point, 75 µl/well were transferred to a new plate and immediately frozen at -80 °C (cells and supernatants), the remaining 75 µl/well were centrifuged for 5 min at 4.680 rpm in an Eppendorf 5430 centrifuge, and 50 µl of the supernatants transferred to a new plate, which was immediately frozen at -80 °C (supernatants). To assess drug stability in the culture media (mGAM), drugs were pooled together in three different pools according to drug-treatment concentration: 80 mM, 20 and 30 mM and 2.5 and 5 mM. Immediately after pooling the drugs, 1) 100 µl were transferred to 96-well shallow U-bottom plates and immediately stored frozen at -80 °C (0 h time point), and 2) 150 µl were transferred to 5 different 96-well U-bottom plates, which were incubated anaerobically at 37 °C for 1.15 h, 2.5 h, 5 h, 7.5 h, and 10 h, respectively. At each time point, the plates were immediately frozen at -80 °C. Metabolite extraction. Frozen samples were thawed on ice, and 20 µl of supernatants, cells and supernatants or of mGAM were distributed into 96-well plates (V-bottom storage plates, Thermo Scientific) according to drug-treatment concentration: 80 mM, 20 and 30 mM and 2.5 and 5 mM. Additionally, 20 µl of pooled drug stocks, split according to the concentration range, were serial diluted in fresh mGAM to set-up calibration curves in all plates. Pooled ^13^C-niclosamide, warfarin, caffeine, iripiflavone, 2-amino-N-cholrophenyl)benzamide (Sigma), and sulfamethoxazole (TOKU-E) were diluted in 50% DMSO and used as internal standards (IS) by adding 5 µl per well of 160 mM, 40 mM and 10 mM stock concentrations to the 80 mM, 20 and 30 mM, and 2.5 and 5 mM sample plates, respectively. For metabolite extraction, 100 µl/well of a 1:1 mix of methanol:acetonitrile were added using the Biomek FXP (Beckman Coulter) liquid handling system. Well contents were mixed and plates were incubated at -20 °C overnight. After incubation, samples were centrifuged at 4.000 rpm in a 5810R eppendorf centrifuge for 10 min at 4 °C. 80 mM, 20 and 30 mM, and 2.5 and 5 mM treated samples were diluted 1/7, 1/2, and 2/1, respectively, with H_2_O for analysis by LC-MS. These experiments were performed in 2 biological replicates with 2 technical replicates each. LC-MS measurements. Chromatographic separation was performed using an Agilent InfinityLab Poroshell HPH-C18 1.9 µM, 2.1 x 10 mm column and an Agilent 1290 Infinity II LC system coupled to a 6546 Q-TOF mass spectrometer. Column temperature was maintained at 45 °C with a flowrate of 0.6 ml/min. The following mobile phases were used: Mobile phase A: Water with 0.1% Formic acid and mobile Phase B: Acetonitrile with 0.1% Formic acid. 5 mL of sample were injected at 5% mobile phase B, maintained for 0.10 min, followed by a linear gradient to 95% B in 5.5 min and maintained at 95% B for 1 min. The column was allowed to re-equilibrate with starting conditions for 0.5 min before each sample injection. The mass spectrometer was operated in positive mode (50–1,700 m/z) with the following source parameters: VCap, 3,500 V; nozzle voltage, 2000 V; gas temperature, 275=°C; drying gas 13 l/min; nebulizer, 40 psi; sheath gas temperature 275=°C; sheath gas flow 12 l/min, fragmentor, 365 V and skimmer, 750 V. Online mass calibration was performed using a second ionization source and a constant flow (10 µL/min) of reference mass solvent (121.0509 and 922.0098 for positive). Quantification of compounds was performed using the MassHunter Quantitative Analysis Softwere (Aglient Technologies, version 10.0). For niclosamide, we used the same method as we reported for single species metabolomics (see below). During data processing, one of the internal standards was selected to normalise the compound’s signal based on the correlation between the measured signal and the expected concentration.

### Chemical screening of bacterial monocultures

#### Screening plates preparation

Drugs were dissolved with the appropriate solvent (Table S1) at a concentration 200-fold higher than the screening concentration, distributed in 96-well V-bottom plates (Greiner, 651261), each well containing 10 µl of dissolved compound or vehicle, and stored at -30°C for up to 1 month. Before the experiment, drug plates were pre-reduced overnight in the anaerobic chamber inside an ice-box at 4 °C. The experiment day, the 10 µl of drug or vehicle was added to 990 µl/well of media in 96-deep well plates (Costar 3959) and 50 µl/well were transferred to shallow U-bottom 96-well plates using the epMotion 96 (Bio-Rad) semi-automated electronic 96 channel pipette.

#### Species inoculation

Species, started in liquid media from a -80 °C glycerol stock, were passaged twice overnight. For inoculation, the second overnight culture was diluted into fresh medium to an OD_578_ of 0.02 (2x). 50 µl/well were added to the 96-well U-bottom drug containing plates, to final drug concentrations as indicated in Table S2, 1% DMSO, and starting bacterial cultures at OD_578_ of 0.01. Plates were sealed with breathable membranes and incubated at 37 °C, with shaking only a few seconds before OD measurement, as described above. These experiments were performed in 3 biological replicates with 2 technical replicates each.

#### Single-species growth curves

Growth curves for single species were determined for the 21 species that were consistently detected in the in vitro communities (Table S1). As previously described ^3^, growth curves were quality-controlled, and truncated at the end of the exponential phase under control conditions. The AUC was calculated for all conditions and divided by the median control AUC within each plate. Across all treatment conditions, the standard deviation between biological and technical replicates had a mean value of 0.04 and 0.03, respectively. In our previous screen ^3^, a large number of control wells and wells with inactive drugs made it possible to calculate a distribution of AUCs for normal growth, and to calculate p-values based on this distribution. In this more focused screen, this was not possible and we therefore opted to use an AUC threshold of 0.75 (i.e. 25% reduced growth) to determine whether a species was susceptible to drug treatment.

#### Single-species metabolomics

LC-MS measurements were employed to assess the presence/absence of a drug or a drug metabolite after incubation with a single-species culture. A 96-deep well plate was filled with 750 µl/well of a 2x drug concentration in mGAM. Inoculation was performed by filling the plate with 750 µl/well of mGAM containing different species at OD_578_ of 0.02, as indicated. Immediately after species inoculation: 1) 100 µl were transferred to 96-well U-bottom plates for growth curve acquisition, as described above, 2) 100 µl were transferred to 96-well U-bottom plates and immediately stored frozen at -80 °C (0 h time point), and 3) 100 µl were transferred to different 96-well U-bottom plates, which were incubated anaerobically at 37 °C for the indicated times. At each time point, plates were immediately frozen and stored at -80°C until metabolite extraction. Metabolite extraction. Frozen samples were thawed on ice, and 20 µl of the samples were distributed into 96-well plates (V-bottom storage plates, Thermo Scientific). Additionally, 20 µl of pooled niclosamide and aminoniclosamide, or of nifurtimox were serial diluted in fresh mGAM to set-up calibration curves. Pooled 2-amino-N-cholrophenyl)benzamide and ^13^C-niclosamide (Sigma) were diluted in 50% DMSO and used as internal standards (IS) by adding 5 µl per well of 2x drug treatment respectively. Metabolite extraction was performed with methanol:acetonitrile as described above. These experiments were performed at least in 2 biological replicates with 2 technical replicates each. When specified, drug bioaccumulation was analyzed by splitting supernatants from cells and supernatants by centrifugation right after time point collection and before freezing at -80 °C, as described above for the community metabolomics. Bioaccumulated drug concentration was calculated by subtracting the drug concentration obtained from the cell and supernatant fraction minus the drug concentration in the supernatant fraction. LC-MS measurements. Chromatographic separation was performed using an Agilent InfinityLab Zorbax Eclipse Plus C18, 1.8mM, 2.1 x 50mm column and an Agilent 1290 Infinity II LC system coupled to a 6546 Q-TOF mass spectrometer. Column temperature was maintained at 45 °C with a flow rate of 0.6 ml/min. The following mobile phases were used: Mobile phase A: Water with 0.1% Formic acid and mobile Phase B: Acetonitrile with 0.1% Formic acid. 5 mL of sample were injected at 5% mobile phase B, maintained for 0.10 min, followed by a linear gradient to 20% B in 0.4 min, followed by a linear gradient to 95% B in 4 min and maintained at 95% B for 0.5 min. The column was allowed to re-equilibrate with starting conditions for 0.5 min before each sample injection. The mass spectrometer was operated in positive mode for initial 3.6 min and then switched to negative scanning mode (50–1,700 m/z) with the following source parameters: VCap, 4,000 V; nozzle voltage, 1000 V; gas temperature, 290=°C; drying gas 13 l/min; nebulizer, 50 psi; sheath gas temperature 400=°C; sheath gas flow 12 l/min, fragmentor, 130 V and skimmer, 750 V. Online mass calibration was performed using a second ionization source and a constant flow (10 µL/min) of reference mass solvent (121.0509 and 922.0098 for positive and 119.0363 and 1033.9881 m/z for negative operation mode, respectively). Quantification of compound was performed using the MassHunter Quantitative Analysis Softwere (Aglient Technologies, version 10.0).

### Quantification of community effects

#### Protection and sensitization

The assessment of the effects of the community on the drug sensitivity of individual species is based on the comparison of the expected behavior in bacterial monocultures and the observed treatment effects in the bacterial community. To compute the fraction of species that are protected in the community for a certain treatment condition, we only considered the subset of species that are affected by the drug in the monoculture experiment. For this subset of species, we divided the number of species that grew normally in the community by the number of affected species. Conversely, to determine the fraction of species that are cross-sensitized in the community we divided the number of species that showed reduced growth in the community (but grew normally in monoculture) by the number of species that grew normally in the monoculture experiment. This calculation was done on the level of technical replicates from the community experiment.

#### Concentration dependency

The concentration dependency of the community effects was calculated both on the concentration steps of the drug (low/middle/high) and the actual numerical concentration. In the first instance, we determined for each drug the concentration steps for which the community effect could be determined. For example, consider a drug which only reduced the growth of any species monoculture at its middle and high concentration but not at its lowest concentrations. For determining the concentration dependency, these would be considered as the low and middle concentration. The distributions of the fractions of affected species were compared using Wilcoxon signed-rank tests.

In addition, we fitted separate logistic functions for sensitization and protection for all drugs which were affected at more than one concentration step. Parameters are a common growth rate k and drug-specific offset d_i_ to capture the relation between the fraction of affected species f_i,j_ for the concentration step c_i,j_:

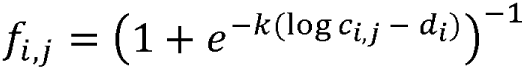

We compared the resulting model with a simplified model that has no concentration dependency, i.e., only one constant value for each drug. We ascertained that the concentration-dependent model provides a better fit to the data using both an ANOVA (analysis of variance) and the BIC (Bayesian information criterion).

### Spent media experiments

Spent media experiments were used to evaluate specie’s effects on drugs and whether these protect (potential protectors) other species (drug-sensitive species) from drug toxicity. All species were inoculated from glycerol stocks into liquid medium, passaged for two days, and diluted to final OD_578_ of 0.01 to perform the experiments. Potential protectors and mGAM controls were grown, for the indicated times, in the presence of 2x the indicated drug concentrations. After the incubation periods, the potential protectors and mGAM controls were sterilized with 0.22 µm PVDF filters (Millipore). 50% of the filtrate (growth spent media) was diluted with 50% of fresh mGAM containing drug-sensitive species at OD_578_ of 0.02 in shallow 96-well U-bottom drug-containing plates. The final drug concentrations are as indicated in the figures. Plates were sealed with breathable membranes and incubated at 37°C for monitoring the growth of the sensitive species every hour during 24 h, as described above. These experiments were performed at least in 3 biological replicates with 2 technical replicates each.

### Small community growth rescue by a drug detoxifier

Two small bacterial communities assembled by mixing *Bacteroides uniformis*, *Bifidobacterium longum*, *Streptococcus parasanguinis*, *S. salivarius* with and without *Coprococcus comes* at a starting OD_578_ nm of 0.01 were kept untreated or treated with 10 µM niclosamide for 24 h. At 24 h pellets were collected by centrifugation and DNA was isolated and bacterial community composition was quantified by qPCR. Community growth was monitored in 96-well U-bottom plates as described above.

#### Bacterial community composition quantification by qPCR

##### Primer design

Species-specific primers were designed using the NCBI primer design tool by selecting the following common parameters: product length of 200 bp maximum, 25 bp primer length, Tm 70 °C ± 3 °C, G/C content < 60%. Primers were tested *in silico* for homology to non-specific sites against nr and RefSeq representative genomes by BLAST, and *in vitro* by PCR against genomic DNA isolated from each individual species included in the community. The 515F/806R primers ^35^ (Table S2) targeting the V4 hypervariable regions of 16S rDNA were used for the amplification of the 16S rDNA of the community in each sample. Four-point standard curves were prepared from ten-fold serial dilutions of DNA prepared for each individual species, starting at concentration of ∼1 ng/µL using 1 µL per well in duplicate reactions (range from ∼1 ng – 2 pg). The standard curves demonstrated good linearity in four orders of magnitude (R^2^ of 0.989-1.000) for all DNAs except for DNA isolated from *B. uniformis*, where linearity was three orders of magnitude. Species specific primer’s efficiencies ranged from 91 to 100 %, 16S rDNA primer’s efficiencies were 84.5% ± 5.5%. *Reaction conditions.* 20 µL qPCR reactions containing 10 µL SYBR^TM^ master mix (Applied Biosystems), 1 µL nuclease-free water, 4 µL of 4 µM primer mix (0.2 µM each final), and 1 µL of template DNA (5-10 ng) were run on 96-well plates on a StepOne Plus Real-Time PCR system (ThermoFisher Scientific). A melting curve analysis was carried out to confirm the amplification of a single product in each reaction. *Data analysis.* Normalized DNA contributed by each species per condition was determined by calculating the threshold cycle (ΔΔCT) value for each gene in relation to the 16S rDNA of the community. To reflect the effect of the community growth on the final community composition, the resulting normalized DNA value was divided by the final OD_578_ of the community at 24 h.

### Overexpression of nitroreductases

#### Proteomics

For proteomics experiments, overnight cultures of the different species were diluted to an OD_578_ 0.1, and anaerobically grown until OD_578_∼1. At this point, cell cultures were treated with vehicle (DMSO) or 20 µM niclosamide for 30 minutes. Cell pellets were collected by 10 min centrifugation at 4000 x g, washed with ice-cold PBS, and frozen at -80°C until further processing. Cells were then resuspended in a lysis buffer (2% SDS, 250 U/ml benzonase, and 1 mM MgCl_2_ in PBS) and boiled at 95 °C for 10 min. After measuring protein concentration using the BCA assay, according to the manufacturer’s instructions (ThermoFisher Scientific), 5 µg of protein from each condition were digested using a modified SP3 protocol ^38^ as previously described ^39^. Peptides were labeled with TMT10plex (Thermo Fisher Scientific), fractionated under high pH conditions, and analyzed using liquid chromatography coupled to tandem mass spectrometry, as previously described ^39^. Mass spectrometry data were processed using isobarQuant and Mascot 2.4 (Matrix Science) against the R. intestinalis (UP000004828) and P. vulgatus (UP000002861) UniProt FASTA. Protein abundance (niclosamide treated samples vs vehicle control) was calculated after normalizing the signal sum intensity across conditions using vsn ^40^.

#### Nitroreductases over-expressing species

Annotated R. intestinalis (UP000004828) and P. vulgatus (UP000002861) nucleotide sequences of nitroreductases were retrieved from the genome annotation databases from UniprotKB ^41^. Plasmids were constructed by Gibson assembly reactions, using the plasmid pww3864 as a backbone ^42^, which were transformed into the chemically competent E. coli species EC100. For this, PCRs using the primer sequences described in Table S2 were used to amplify the vector pww3864 and the nitroreductases ORFs from extracted genomic DNA. Resulting plasmids and pww3864 were electroporated into the E. coli conjugation donor species EC100D pir+ RP4-2-Tc::[ΔMu1::aac(3)IV-ΔaphA-Δnic35-ΔMu2::zeo] ΔdapA::hygromycin. For conjugating the nitroreductases plasmids from the E. coli donor into the recipient P. vulgatus, overnight aerobic cultures of donor species (grown in LB + 0.3 mM diaminopimelic acid (DAP) + 100 µg/ml ampicillin) and anaerobic cultures of P. vulgatus (grown in mGAM) were diluted 50- and 20-fold respectively, and grown for 3 h. At 3 h, cells were collected by centrifugation and donor cells were washed 3 times with LB. Cells were resuspended in 200 µl of mGAM, and 25 µl of each donor species was mixed with 25 µl of the recipient P. vulgatus, plated on mGAM plates supplemented with 0.3 mM DAP, and incubated at 37 °C overnight aerobically. After overnight incubation, cells were collected and plated on mGAM plates supplemented with erythromycin (10 µg/ml) and gentamicin (200 µg/ml). P. vulgatus transconjugants were confirmed by PCR.

## Data availability

Data is available from https://github.com/grp-bork/drugbug_Santamarina_2023.

Raw 16S amplicon sequencing data has been deposited at the European Nucleotide Archive (ENA) under accession number PRJEB46619: https://www.ebi.ac.uk/ena/browser/view/PRJEB63118.

## Code availability

Code is available at https://github.com/grp-bork/drugbug_Santamarina_2023

## Supplemental information

### Supplementary tables

**Supplementary Table 1**

List of gut bacterial species, drugs, colon concentrations, low abundant species, a single-species AUCs.

**Supplementary Table 2**

Putative FMN-dependent nitroreductases in the gut bacterial species from this study, differential proteomics analysis of *R. intestinalis* and *P. vulgatus* nitroreductases overexpressing strains upon niclosamide treatment, and list of primers used in this study.

## Notes

https://github.com/grp-bork/drugbug_Santamarina_2023

https://www.ebi.ac.uk/ena/browser/view/PRJEB63118

### Competing Interest Statement

The authors have declared no competing interest.

