## Supplemental Figures for "Emergence of community behaviors in the gut microbiota upon drug treatment"

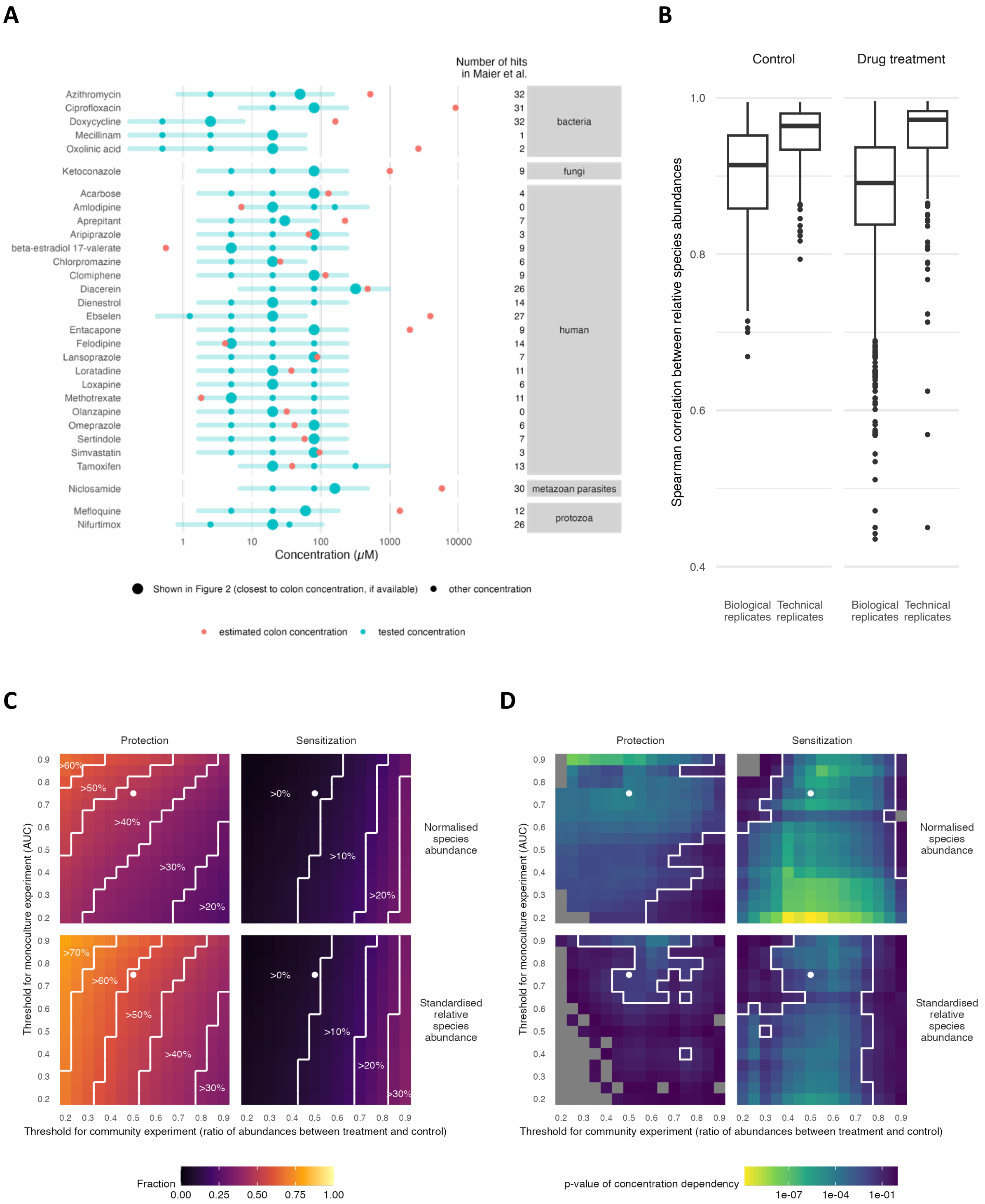
A
Number of hits in Maier et al.
Azithromycin 32
Ciprofloxacin 31
Doxycycline 32
Mecillinam 1
Oxolinic acid 2
bacteria
Ketoconazole 9
fungi
Acarbose 4
Amlodipine 0
Aprepitant 7
Aripiprazole 3
beta-estradiol 17-valerate 9
Chlorpromazine 6
Clomiphene 9
Diacerein 26
Dienestrol 14
Ebselen 27
Entacapone 9
Felodipine 14
Lansoprazole 7
Loratadine 11
Loxapine 6
Methotrexate 11
Olanzapine 0
Omeprazole 6
Sertindole 7
Simvastatin 3
Tamoxifen 13
human
Niclosamide 30
metazoan parasites
Mefloquine 12
Nifurtimox 26
protozoa
1
10
100
1000
10000
Concentration (μM)
Shown in Figure 2 (closest to colon concentration, if available)
other concentration
estimated colon concentration
tested concentration
B
Control
Drug treatment
Spearman correlation between relative species abundances
1.0
0.8
0.6
0.4
Biological replicates
Technical replicates
Biological replicates
Technical replicates
C
Protection
Sensitization
>60%
>50%
>40%
>30%
>20%
>0%
>10%
>20%
Normalised species abundance
>70%
>60%
>50%
>40%
>30%
>0%
>10%
>20%
>30%
Standardised relative species abundance
Threshold for monoculture experiment (AUC)
Threshold for community experiment (ratio of abundances between treatment and control)
0.2 0.3 0.4 0.5 0.6 0.7 0.8 0.9
Fraction
0.00 0.25 0.50 0.75 1.00
D
Protection
Sensitization
Normalised species abundance
Standardised relative species abundance
Threshold for monoculture experiment (AUC)
Threshold for community experiment (ratio of abundances between treatment and control)
0.2 0.3 0.4 0.5 0.6 0.7 0.8 0.9
p-value of concentration dependency
1e-07 1e-04 1e-01

**Suppl. Fig. 1. Drug characteristics, quality control and threshold exploration.**
(**A**) The drug concentrations tested in the screen. Selection was based on prior knowledge of drugs' activity at 20 µM [1] (counts of species targeted on the right side). Estimated colon concentrations are shown as a red dot (when available), and tested concertation closest to the estimated colon concentration are show as enlarged dots (if colon concentration not available, then the 20 µM is shown).

(**B**) Distribution of Spearman correlations between relative abundances for technical and biological replicates for 16S sequencing.

(**C**) Exploration of thresholds for defining growth inhibition in the community and in monoculture. Cells are colored by the fraction of protected (left) or sensitized (right) species for pairs of thresholds. The white dot designates the chosen thresholds of 25% growth inhibition in monoculture and 50% growth inhibition in the community. White lines delineate regions of similar fractions. The fractions of protection and sensitization changed only slowly as the cutoffs for determining growth inhibition were varied. Likewise, the abundance normalization based on OD produces similar results as the standardization of relative abundances by dividing by the 75$^{th}$ percentile of relative abundances.

(**D**) The concentration dependency of the fraction of protection and sensitization is not sensitive to the chosen thresholds of defining growth inhibition. The white line encloses the region where the p-value of the concentration dependency is below 0.05. Grey cells are shown for threshold pairs where the sigmoid curve fitting for the concentration dependency failed.

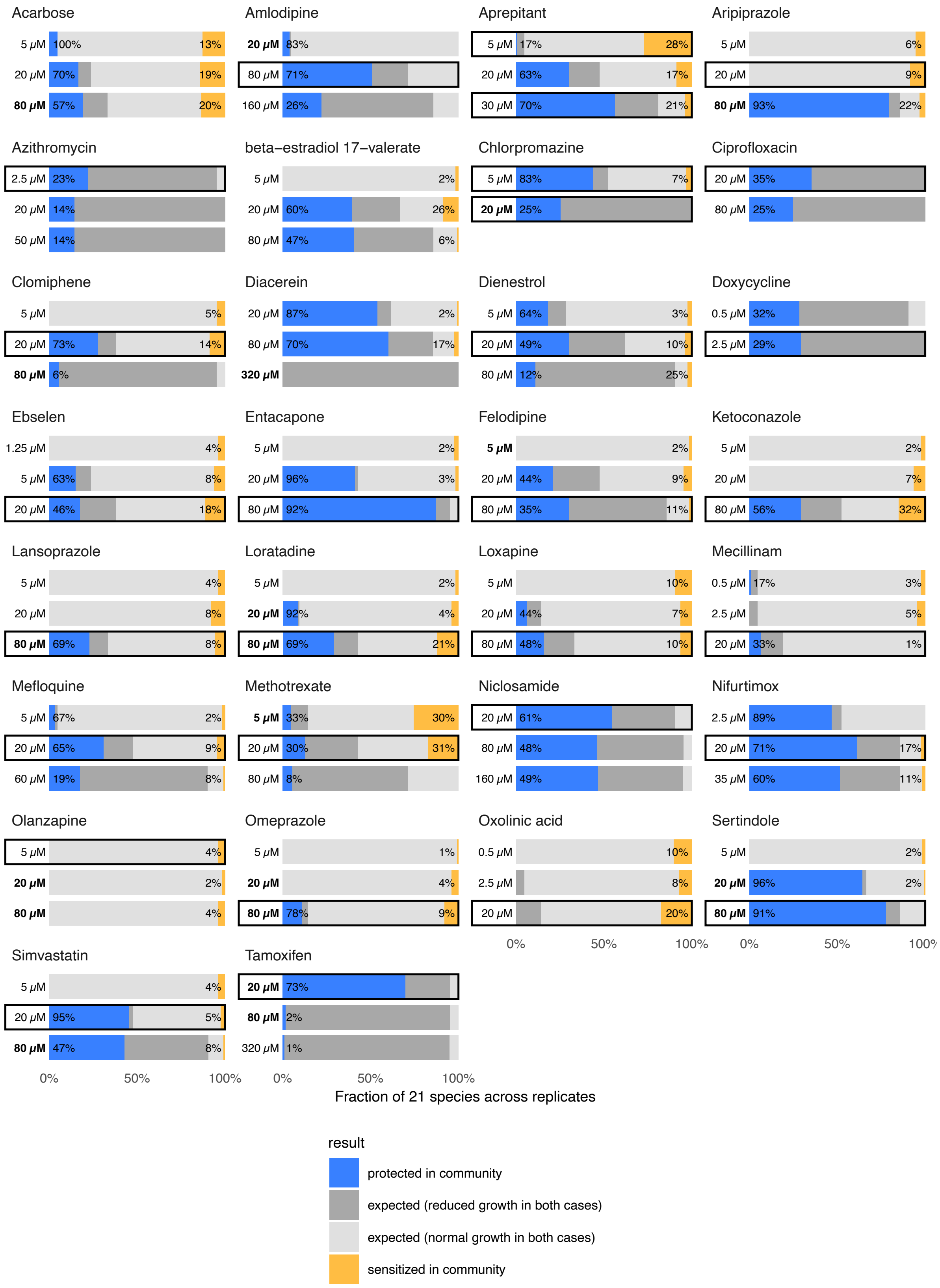


**Suppl. Figure 2. Overview of the emergent behaviors for all drugs and concentration steps**. For each treatment condition, the range of community effects is shown. Percentages of protection are shown relative to species being inhibited in monoculture (blue/blue + dark grey), and percentages of cross-sensitization are shown relative to species being unaffected in monoculture (yellow/yellow + light grey). Concentrations in bold: Within a factor of three of the estimated gut concentration. Outlined in rectangles: Conditions chosen for follow-up metabolomics experiments.

A

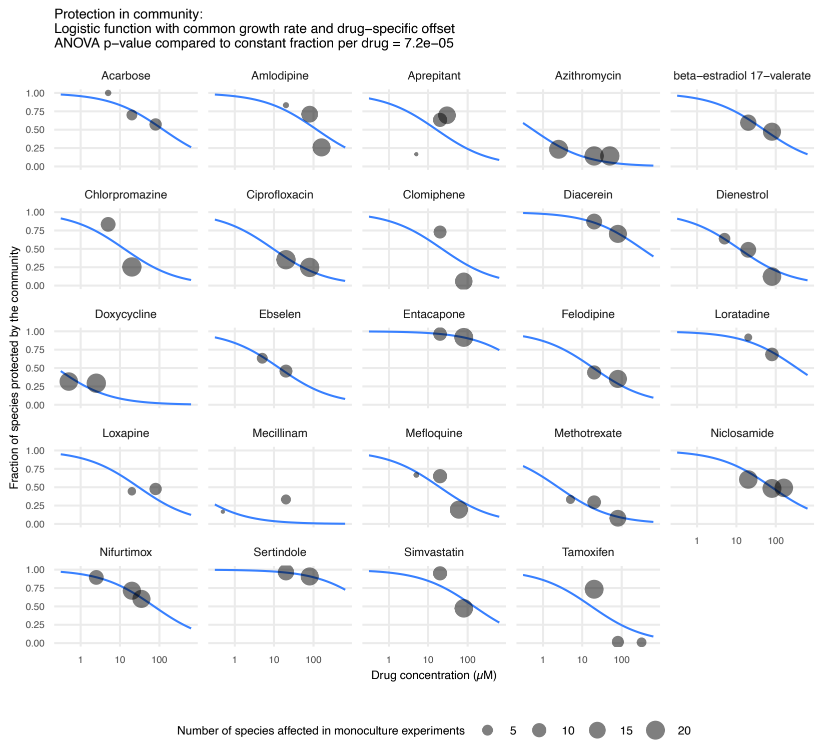


B

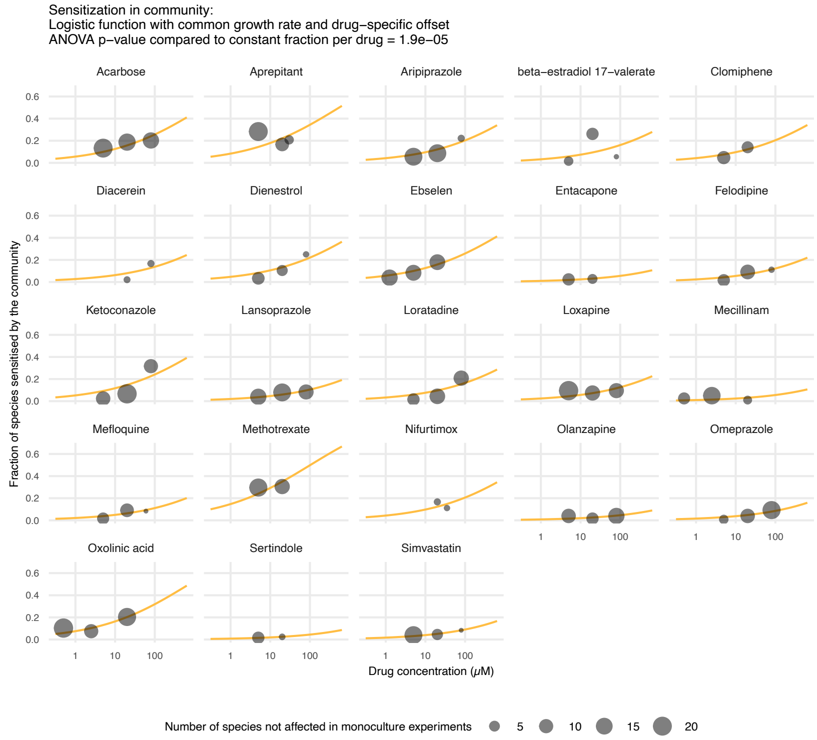


**Suppl. Figure 3. Concentration dependency of emergent behaviors.**

(**A-B**) The fraction of protected species is shown as a function of the drug concentration. We modeled the dependency between the drug concentration and the fraction of affected species as a logistic function, which has two parameters: the growth rate and the offset (i.e. the concentration where the fraction of the affected species would be 0.5). For the curve fitting, we used the same growth rate for all drugs, but drug-specific offsets. Hence, the "shape" of the logistic curve is the same across all drugs, but the curve is shifted according to the drugs' overall effect on the community. An ANOVA was used to compare the logistic function fit to a concentration-independent model with a constant fraction of affected species per drug. B is same as (A), but this time for cross-sensitization.

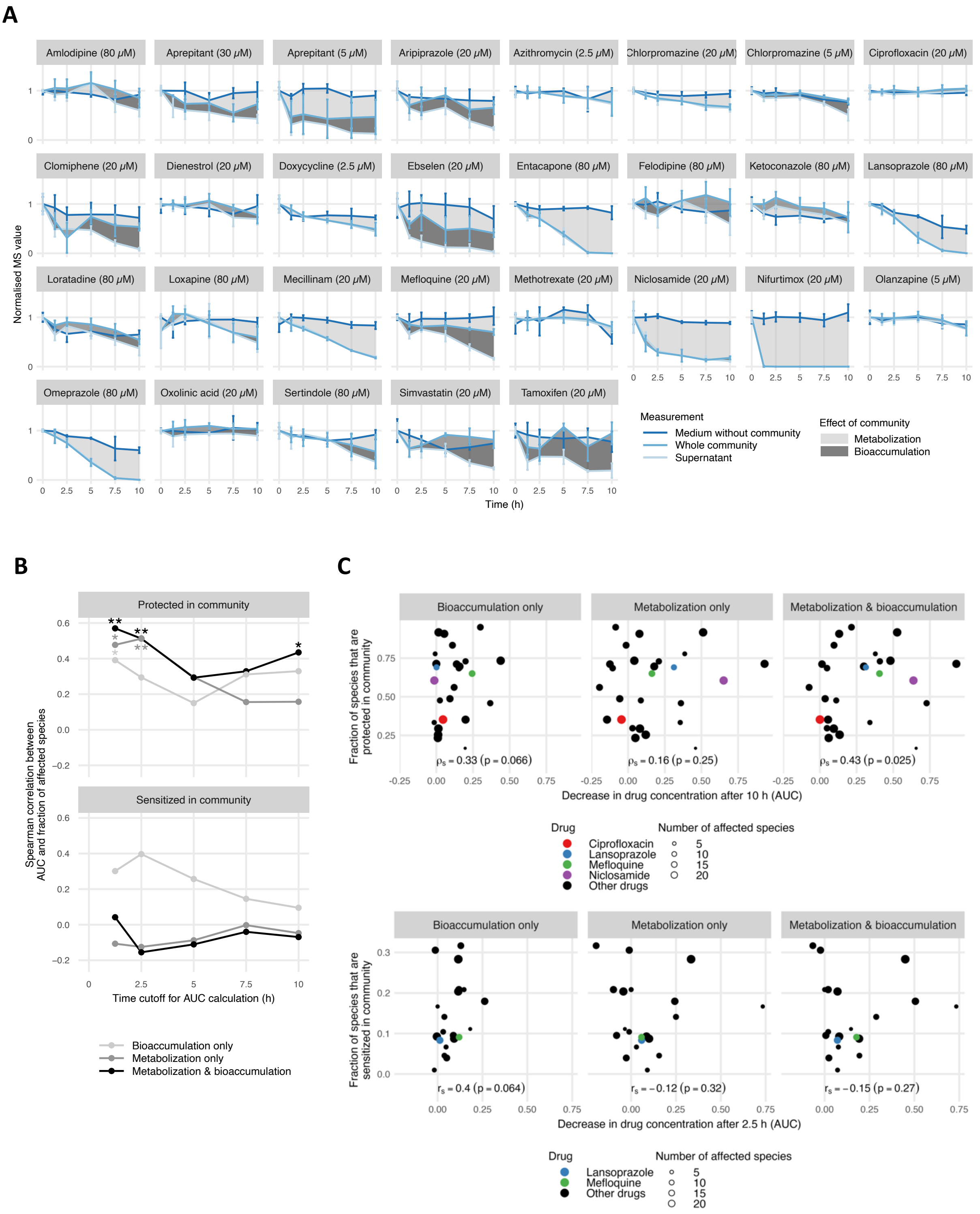


**Suppl. Figure 4. Metabolomics profiling of communities explains large part of cross-protection.**

(**A**) Time-course of biotransformation and bioaccumulation for all drugs (graphs plotted as in Fig. 3A)

(**B)** The correlation between the compound biotransformation and/or bioaccumulation and the fraction of protected (top) and sensitized (bottom) species as a function of different time cutoffs in the metabolomics data. Asterisks: ** : $p < 0.01$, * : $p < 0.05$.

(**C**) Fraction of sensitized species in the community as a function of drug biotransformation and/or bioaccumulation at 2.5 hr. Only bioaccumulation shows some correlation, albeit it fails to reach significance. P-values were calculated by permutation tests using 100,000 samples.

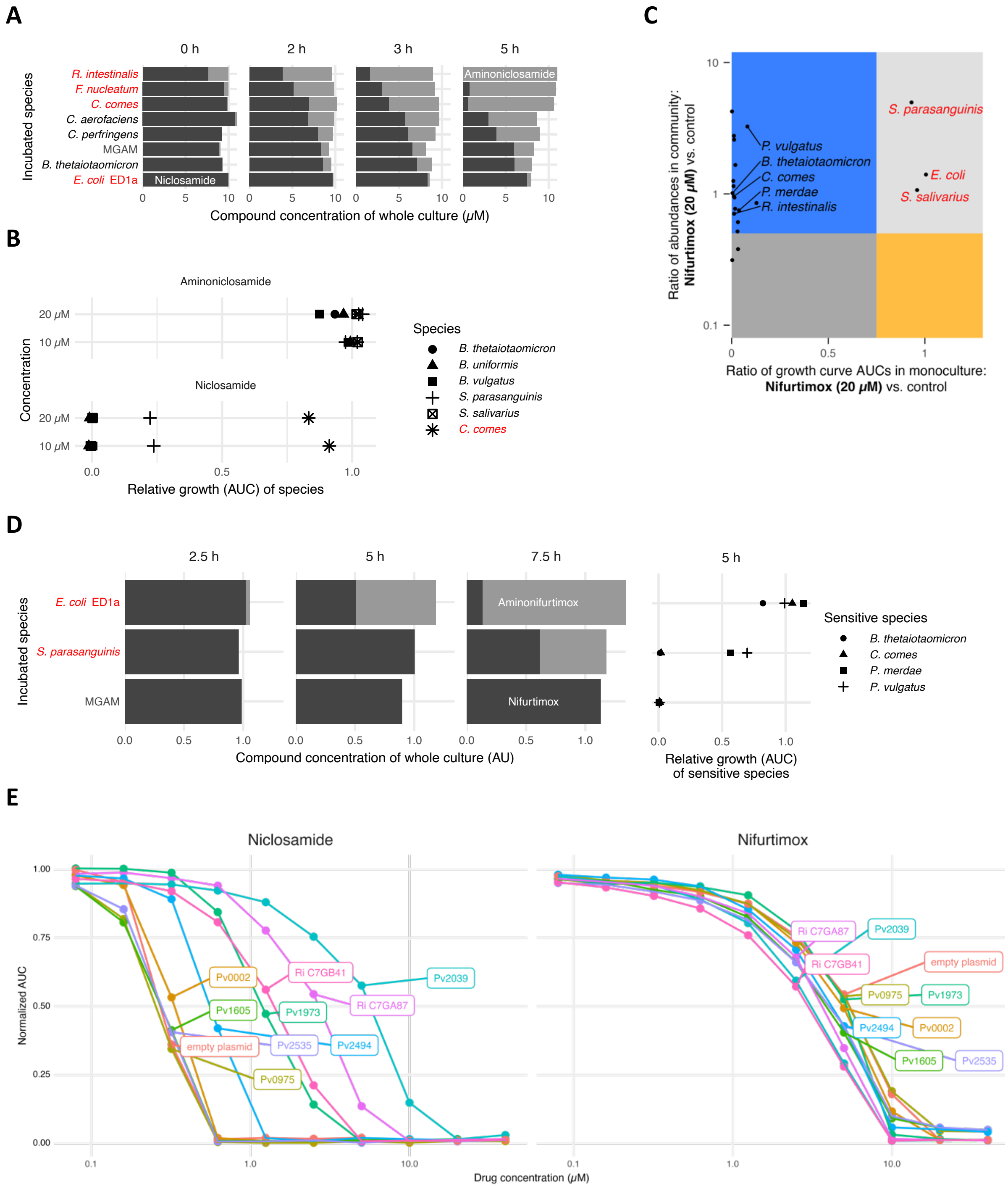


**Suppl. Figure 5. Contribution of individual strains to niclosamide and nifurtimox community protection.**

(**A**) Full time-course for the niclosamide degradation experiment shown in Fig. 4B.

(**B**) Relative growth of niclosamide-sensitive (black) and resistant (red) species subjected to niclosamide (control) and aminoniclosamide treatments. Aminoniclosamide was non-toxic to all species.

(**C**) Single-species versus community growth phenotypes upon treatment with 20 µM nifurtimox. Representation of species that grow in community as expected from monoculture (light and dark grey areas), or that are cross-protected (blue area) or cross-sensitized (yellow area) in the community.

**(D)** Full time-course for nifurtimox biotransformation (as in **A**). In contrast to niclosamide, *E. coli* strongly reduced nifurtimox to aminonifurtimox. Spent media from *E. coli* and *S. parasanguinis* (resistant species to Nifurtimox – panel **C**), grown in the presence of 40 µM nifurtimox (or vehicle) for 5 hours, was mixed 1:1 with fresh mGAM, and used to grow nifurtimox sensitive strains. Growth of the sensitive was normalized to their growth in spent media from untreated cultures.

(**E**) Niclosamide and nifurtimox dose response curves for *P. vulgatus* heterologously overexpressing different nitroreductases. Although specific nitroreductases increased the dose response and $IC_{90}$ to niclosamide (Fig. 5A), minor or no changes were observed for nifurtimox.

1. Maier, L., Pruteanu, M., Kuhn, M., Zeller, G., Telzerow, A., Anderson, E.E., Brochado, A.R., Fernandez, K.C., Dose, H., Mori, H., et al. (2018). Extensive impact of non-antibiotic drugs on human gut bacteria. Nature *555*, 623–628. 10.1038/nature25979.
